# Gray whale transcriptome reveals longevity adaptations associated with DNA repair and ubiquitination

**DOI:** 10.1101/754218

**Authors:** Dmitri Toren, Anton Kulaga, Mineshbhai Jethva, Eitan Rubin, Anastasia V Snezhkina, Anna V Kudryavtseva, Dmitry Nowicki, Robi Tacutu, Alexey A Moskalev, Vadim E Fraifeld

## Abstract

One important question in aging research is how differences in genomics and transcriptomics determine the maximum lifespan in various species. Despite recent progress, much is still unclear on the topic, partly due to the lack of samples in non-model organisms and due to challenges in direct comparisons of transcriptomes from different species. The novel ranking-based method that we employ here is used to analyze gene expression in the gray whale and compare its *de novo* assembled transcriptome with that of other long- and short-lived mammals. Gray whales are among the top 1% longest-lived mammals. Despite the extreme environment, or maybe due to a remarkable adaptation to its habitat (intermittent hypoxia, Arctic water, and high pressure), gray whales reach at least the age of 77 years. In this work, we show that long-lived mammals share common gene expression patterns between themselves, including high expression of DNA maintenance and repair, ubiquitination, apoptosis, and immune responses. Additionally, the level of expression for gray whale orthologs of pro- and anti-longevity genes found in model organisms is in support of their alleged role and direction in lifespan determination. Remarkably, among highly expressed pro-longevity genes many are stress-related, reflecting an adaptation to extreme environmental conditions. The conducted analysis suggests that the gray whale potentially possesses high resistance to cancer and stress, at least in part ensuring its longevity. This new transcriptome assembly also provides important resources to support the efforts of maintaining the endangered population of gray whales.

## INTRODUCTION

Long-lived animals represent a unique model for investigating the evolution of longevity. The recently conducted analyses of the longest-lived mammal, studying the bowhead whale (*Balaena mysticetus*) transcriptome (Seim et al., 2014) and genome (Keane et al., 2015), showed that genetic and transcriptional patterns could, up to a certain extent, explain its extraordinary longevity and resistance to cancer and other age-related diseases. The gray whale (*Eschrichtius robustus*) is also among the top 1% longest-lived mammals. It ranks 8^th^ out of 1012 mammalian species with a known maximum lifespan (Tacutu et al., 2018). Despite the extreme environment, or maybe due to a remarkable adaptation to its habitat (intermittent hypoxia, cold Arctic water, and high pressure), gray whales reach at least the age of 77 years, according to currently available data (Tacutu et al., 2018).

The *Eschrichtius robustus* is the only member of the *Eschrichtiidae* family from the order Cetacea (Wolman, 1985). It is considered a “living fossil” because of its short, coarse baleen plates and lack of a dorsal fin (Nollman, 1999). Gray whales were almost extinct in the middle of the 20th century and despite being protected by law (limited hunting for food being permitted only for the indigenous population of Chukotka) they are still considered to be an endangered species (Weller, Burdin, Wursig, Taylor, & Brownell Jr, 2002). As such, having the opportunity to get insights from its transcriptome is of great importance to aging research.

Here, we comprehensively analyzed the transcriptome of the gray whale in two tissues (liver and kidney), focusing on the possible links between longevity and the expression of individual genes or group of genes. For this purpose, we also compared the gray whale transcriptome (A. A. Moskalev et al., 2017) with that of two other whales, bowhead whale (*Balaena mysticetus*) and minke whale (*Balaenoptera acutorostrata*), and with the transcriptomes of other five mammalian species of different longevities.

## RESULTS AND DISCUSSION

### The gray whale transcriptome assembly and annotation

Kidney and liver mRNA from a young-adult female whale were sequenced using the MiSeq System Illumina sequencer. *De novo* transcriptomes were assembled, yielding in total 114,233 contigs. Orthologous protein-coding genes encoded in the transcriptome were identified using the Sprot (Galgonek, Hoksza, & Skopal, 2011) algorithm and BLASTing (Johnson et al., 2008) against UniProt (Apweiler et al., 2004) sequences. In total, we identified 12,072 protein-coding genes that have been aligned to 7,997 Uniref90 clusters (groups of proteins that have 90% identity). This implies that each of the proteins found by us has homologues in at least one or more mammalian species. Of these, 11,456 are expressed in the liver and 8,363 in the kidney, with a high overlap between tissues (64% of the total). In addition to the identified genes, a large number of the unidentified contigs also fit some of the required criteria for being coding sequences (certain length and expression level). For the full list of unknown genes, please see Dataset S1.

### Enrichment of top-most expressed genes in the gray whale

To understand which gene subsets are most expressed in both tissues and/or maybe representative for the most active processes in an organism, we next conducted a Gene Ontology (GO) enrichment analysis of the top 100 genes (threshold selection described in Experimental Procedures) with the highest contig count compared to all genes (background). Please see Dataset S2 for the list of all contigs and Dataset S3 for the enrichment results. Several interesting features were observed for both liver and kidney. First, we found a high presence of GO categories related to the extracellular matrix (ECM), cell-cell/cell-ECM interactions and exosomes. Interestingly, this is in line with our previous analysis, which highlighted the potential importance of these categories (specifically, focal adhesion and adherens junction proteins) in lifespan determination and in linking human longevity and age-related diseases (Wolfson, Budovsky, Tacutu, & Fraifeld, 2009). Second, the top-most expressed genes were significantly enriched in categories related to immuno-inflammatory responses, organ regeneration and regulation of cell proliferation. One possible explanation might come from the need of large and especially long-lived species like the gray whale to develop strong tumor suppressor mechanisms, in order to compensate for having more somatic cells, and hence a higher risk to develop cancer. The immune system, as well as mechanisms of organ regeneration and regulation of cell proliferation, are crucial for the prevention of cell transformation and elimination of cancer cells (Seluanov, Gladyshev, Vijg, & Gorbunova, 2018). Finally, as expected, based on observations in other species examined so far (Mercer et al., 2011), many top-expressed genes in the gray whale fall into the mitochondria-associated categories. The full list of enriched categories is presented in Dataset S3 for liver and kidney, respectively.

### Unknown genes

Aside from the 12,072 protein-coding genes identified in the gray whale transcriptome, 35% of all assembled contigs remained unidentified. Overall, this percentage of unannotated sequences is not unexpectedly large in the case of *de novo* transcriptome assembly (Keane et al., 2015; Seim et al., 2014). Generally, unidentified sequences might be the result of (i) mapping errors or (ii) still uncharacterized sequences. To further investigate unannotated sequences, while excluding false positives due to mapping, we next selected only those contigs with a high-count number, hence highly expressed, and with a sequence length >200 bp, comparable to that of a common mRNA size (see Experimental Procedures). Out of the 10,389 un-annotated sequences the top 600 have an increased expression, i.e. 10x more than the average in the transcriptome. Among these 600 unannotated, 4 unknown sequences are among the top 100 expressed genes, and in the top 1,000 there are 92 unknown genes, suggesting that their functional significance should be further investigated (Dataset S1).

To look closer at these unannotated sequences, we chose the 20 top-expressed transcripts and manually BLASTed them (Johnson et al., 2008) against the NCBI RefSeq nucleotide database (Haft et al., 2018), searching for similar sequences/domains/motifs (Table S1). Interestingly, all these transcripts were also found highly expressed in other transcriptomes, including those of the bowhead whale and minke whale (data not shown). Swiss-Prot (Apweiler et al., 2004), Pfam (El-Gebali et al., 2018) and the MEME Suite (Bailey et al., 2009) tools did not reveal any significant domain or motif matches in the top-expressed unannotated sequences. Several sequences of predicted/hypothesized proteins or transcripts with high similarity were however found by BLAST in other species. Among them are genes hypothetically related to ribosomal RNA, metabolic processes (Zinc fingers), ERK signaling and cytoskeleton remodeling (ACTB), detoxification of reactive oxygen species, ubiquitination, proliferation, differentiation, and carcinogenesis (PRDX3, CALR), immune response (histocompatibility complex class II and extended class II sub-region, NDRG1, PTMA, GPS2), mitochondria, ATP metabolism and iron binding (PAH, CYP1A1), senescence and autophagy in cancer (CREG1, CAAX box protein), lipoprotein and cholesterol metabolic process (APOA2), regulation of cell differentiation, serpin family (SERPINF2). Interestingly, many of the above-mentioned genes are involved in aging associated pathways (Ma & Gladyshev, 2017; A. Moskalev, Aliper, Smit-McBride, Buzdin, & Zhavoronkov, 2014; Tian, Seluanov, & Gorbunova, 2017).

A broader question is whether some of these unannotated genes are essential for basic/fundamental biological processes. In this regard, Hutchison et al. have recently synthesized a functionally viable artificial genome containing only 473 essential genes. Of them, almost a third (149 genes) had no known biological functions (Hutchison et al., 2016). The fact that unknown genes still exist even in an extremely reduced genome supports our suggestion that many unannotated genes found in the gray whale transcriptome should be a further investigation as they might be involved in fundamental processes.

### Novel approach: Comparative transcriptome analysis of gray whale vs other mammals

To identify expression patterns that are species-specific or shared across species, we then undertook a comparative analysis of the transcriptomes of the gray whale and of seven other mammalian species, including two other whales (bowhead and minke whales) and five terrestrial mammals (naked mole-rat, Brandt’s bat, mouse, cow, and humans) (Fig. 1). These species were selected because (i) the bowhead and minke whales are genetically close to the gray whale, and bowhead whale is the longest-lived mammal (according to existing records); (ii) the naked mole-rat and Brandt’s bat arguably present characteristics of “negligible senescence” and has an impressive lifespan of almost 8 times longer than rodents with comparable body size; (iii) humans are the second longest-lived mammal (according to current records) and are considered among species with exceptional longevity; (iii) the mouse is a good reference for a short-lived organism; and (iv) the cow is a species with a maximum lifespan that is close to the average within the mammalian class (Tacutu et al., 2018) and the most evolutionarily related to whales among modern land mammals (Nikaido, Rooney, & Okada, 1999).

**Fig. 1.**
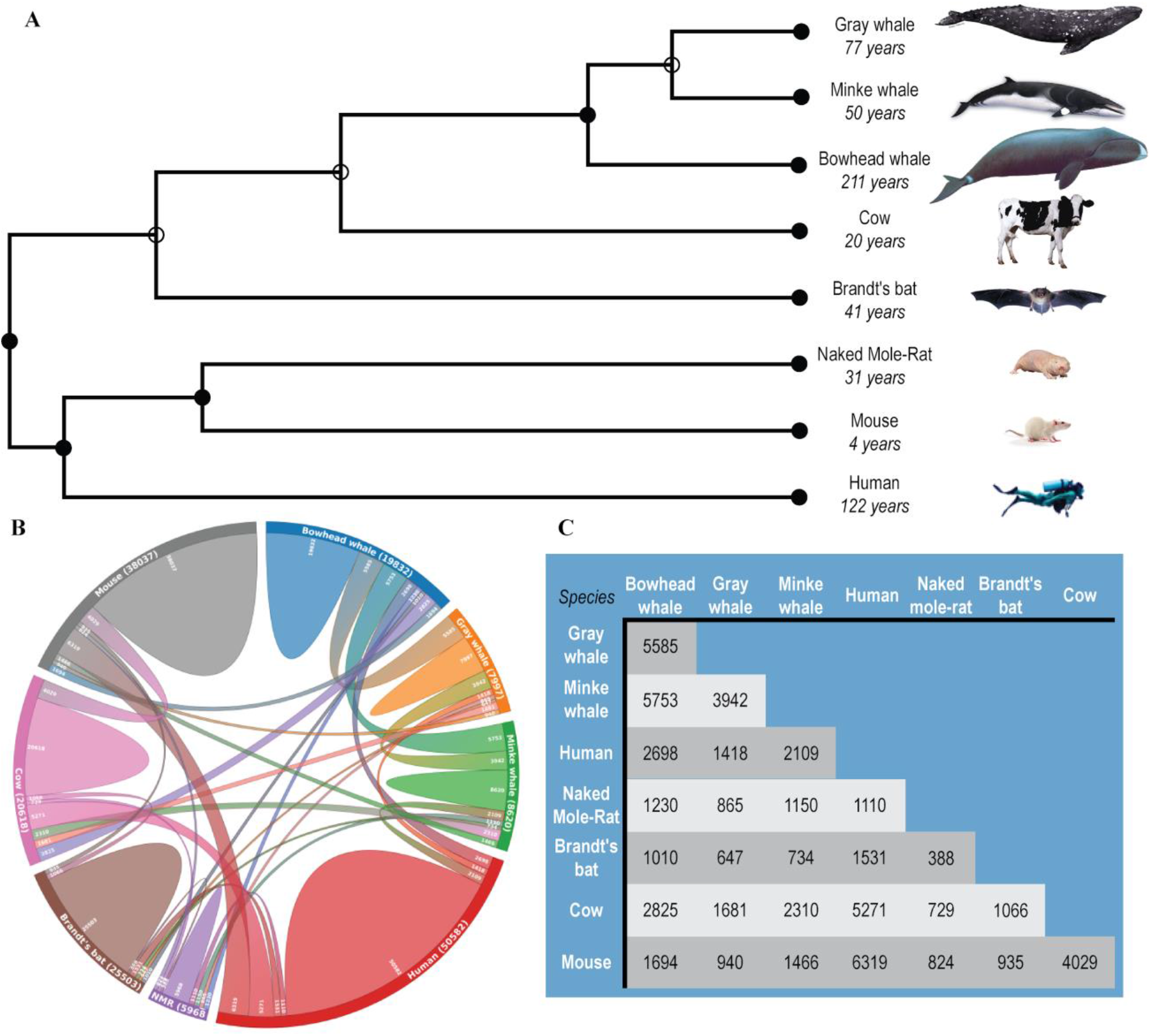
Relationship of the gray whale to other mammalian species. **(A)** Phylogenetic tree of species selected for analysis. The tree with the highest log-likelihood is shown. The percentage of trees in which the associated taxa clustered together (bootstrap values) is shown next to the branches. **(B)** Overlap of Uniref90 protein clusters between the examined species. Uniref90 clusters (which contain proteins with 90% sequence similarity) were predicted from open reading frames (ORFs) extracted from the coding transcripts. The Gray whale transcriptome was assembled in the current study (please see methods), while transcriptomes for other species were taken from previous studies, publicly available at the NCBI database. Presented in brackets is the number of Uniref90 entries predicted from each transcriptome. **(C)** Overlap of Uniref90 protein clusters between the examined species in table format.

Unfortunately, expression levels for individual genes from transcriptomes of different species cannot be compared directly. Additionally, both absolute and normalized expression values could be affected by technical issues. The main culprits are generally the heterogeneity in sequencing methods and/or sequencing equipment, variation in the sample preparation protocols, potential errors in alignments and transcriptome assemblies (Su et al., 2014). To minimize these biases, in our analysis: (i) comparison of species was performed at the level of groups of functionally-linked genes (e.g., GO categories - biological processes) instead of comparing individual genes directly; (ii) within each species, rankings of TPMs were used. For this purpose, the protein-coding transcripts were grouped by GOs and their normalized counts were computed within each GO term. The counts were then ranked, from 1 (the GO with highest composite-expression) to 9779 (the GO with lowest composite-expression) in the gray whale’s transcriptome. Rank values for all species examined are presented in Dataset S4.

To validate the ranking-based approach we compared the intra-species variance in the ranks of a given GO term to the inter-species variance. As expected, the intra-species variance was much lower than the inter-species variance. Phylogenetically closely related species (in our case, gray, bowhead and minke whales) had more similar ranks of a given GO term than more distant species (specifically, whales, on the one hand, and mice and cows, on the other hand). The Spearman correlation analysis indicated a higher similarity between GO transcription level ranks in the same species or closely related whale species (Fig. S1). The values of Spearman’s rank correlation coefficients were 0.97 between the two mice experiments, 0.98 between cow experiments, and ~0.85 for the whales’ category, with a high statistical significance (p < E-25) for both liver and kidney tissues. As expected, the correlation coefficients between the ranks of the gray whale and those of other mammalian species (naked mole-rat, humans, mouse, cow) were much lower (~0.6, p < E-25). These results provide indirectly (however, rather strong) evidence for the validity of GO rank values for comparison of transcriptomes.

Since whales are among the longest-lived mammals, we specifically analyzed the shared processes that are highly expressed in long-lived species, while being lowly expressed in shorter-lived mammals. Our analysis of ranks showed that all longed-lived mammals (whales, humans, naked mole rats, Brandt’s bat) correlate positively between them and negatively with short-lived species (cows and mice) in the following GO categories: DNA maintenance and repair (Fig. 2 A1, A2), autophagy (Fig. 2 B1, B2), ubiquitination (Fig. 2 C1, C2), and to some degree apoptosis (GO categories: “Positive regulation of intrinsic apoptotic signaling pathway in response to DNA damage”, “Endonucleolytic cleavage”, “Positive regulation of endodeoxyribonuclease activity”, “Positive regulation of cysteine-type endopeptidase activity involved in the execution phase of apoptosis”) (Dataset S4). Also, a high similarity between long-lived species was found in the expression of immune response-related GO terms (“Positive regulation of interleukin-2 production”, “Positive regulation of T cell receptor signaling pathway”, “Positive regulation of activated T cell proliferation”). This is in line with previous findings, with a high expression of genes associated with the immune response being recently reported for the bowhead whale transcriptome (Keane et al., 2015; Seim et al., 2014).

**Fig. 2.**
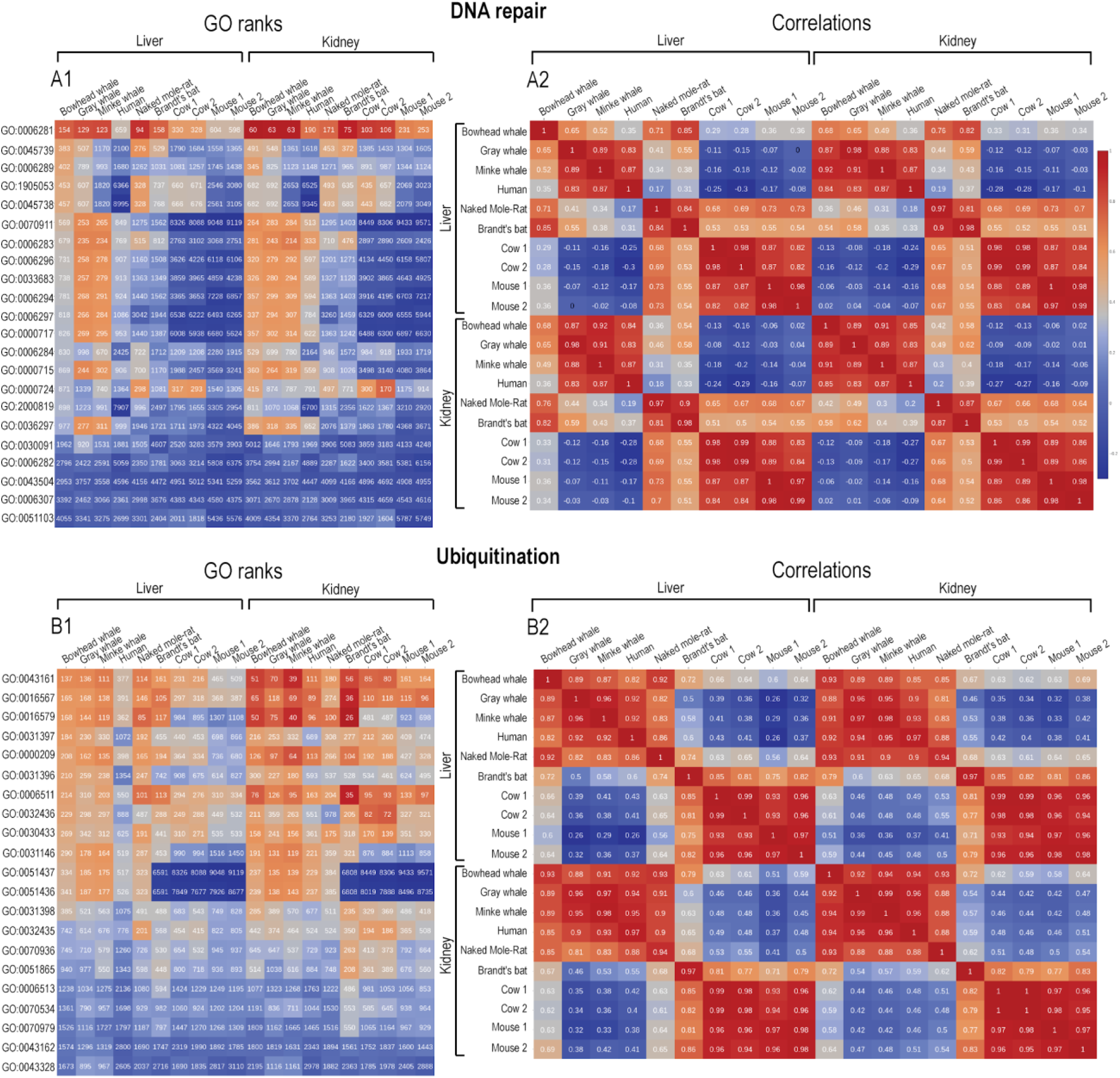
Heatmap of cross-species transcriptome comparative analysis for GO terms. Presented in the figure are the following categories: **A. DNA repair; B. Ubiquitination.** Both liver and kidney tissues are included for all the eight species compared. The same rank ranges and normalization (transcripts per million reads, TPMs) are used for all species. **A1, B1:** Ranks of the top 1,000 expressed GO categories. Ranks range between 1 (top expressed GO term) and 6,260 (least expressed GO term). In the color scheme gradients of red indicate highest expression, gray - middle expression, and blue - lowest expression. A list of names for the GO terms shown in the figure is available in Dataset S5. **A2, B2:** Correlations of GO ranks between every two species. Spearman’s coefficient is shown in the cells of the heatmap. The color gradient indicates the correlation level, from red (highest correlation) to blue (lowest correlation).

Notably, high expression ranks for DNA maintenance and repair, autophagy, ubiquitination (Fig. 2A-C), apoptosis, positive regulation of T cell receptor signaling pathway and activated T cell proliferation (Dataset S4) were also observed for the naked mole-rat, a rodent species with exceptional longevity and remarkable resistance to cancer (Tian et al., 2013). In contrast, in the mouse and cow transcriptome, the expression ranks of the aforementioned categories were much lower, whereas, in humans, the ranks of most GO terms were closer to whales and naked mole rats (Fig. 2A-C; Dataset S4).

The above-mentioned processes are well known to be involved in anti-cancer mechanisms, for preventing cell transformation or elimination of potentially carcinogenic or cancer cells (Cuervo et al., 2005; Seluanov et al., 2018), and in a larger sense in determining mammalian longevity (Keane et al., 2015; Kevei & Hoppe, 2014; Li & de Magalhães, 2013; Yanai, Budovsky, Barzilay, Tacutu, & Fraifeld, 2017). This is highly relevant because while the bowhead whale is known for its resistance to cancer (Seluanov et al., 2018; Tian et al., 2017), for the gray whale, such data is still absent. In light of our findings on the similarity in the expression of anti-cancer processes, it would probably be reasonable to expect that the gray whale might also be resistant to cancer. Additional to the anti-cancer similarities, correlation analysis of expression ranks revealed some interesting patterns for certain GO groups when comparing long-lived with short-lived species. For instance, expression ranks of DNA repair processes show a positive correlation within the groups of long-lived (whales, humans, bats, naked mole rats; p < 5E-4) and short-lived (cows and mice, 2 transcriptomes each; p < 5E-4) species, respectively (Fig. 2 A2). Complementarily, there is a negative correlation for expression ranks between long- and short-lived species (p < 5E-4). A similar result can also be observed for autophagy, ubiquitination (Fig. 2 A2, B2, C2) and other processes like apoptosis and immune response (Dataset S4). Altogether, considering the above high correlations of longevity-associated processes in the gray whale and other long-lived species, the currently accepted record for the gray whale maximum lifespan of 77 years might be an underestimate, stemming from limited available information.

Apart from a high similarity of overall transcription patterns with bowhead and minke whales, the gray whale exhibits a number of particularly high expression ranks for certain gene categories, including GOs relevant to several longevity-related processes. Among them are GO categories like ATP synthesis coupled proton transport, cilia-related processes, regulation of autophagosome assembly, immune responses (the latter two being much higher even than in the bowhead whale), regulation of Wnt signaling pathway, the sensory processes related to the inner ear, cardiac muscle cell differentiation, and neural precursor cell proliferation (see Dataset S4).

### Analysis of longevity-associated genes (LAGs) in the gray whale’s transcriptome

With regards to longevity determination, hundreds of genes have been identified to have an impact, when genetically manipulated, on the lifespan of model organisms like yeast, worm, fruit fly and mouse. These genes have been previously defined as longevity-associated genes (LAGs) and include two categories: a) LAGs which promote longevity, and b) LAGs which reduce lifespan and promote an accelerated aging phenotype (a phenotype that along with shorter lifespan includes features of premature aging) (Budovsky, Abramovich, Cohen, Chalifa-Caspi, & Fraifeld, 2007; Yanai et al., 2017).

Although intuitive, the hypothesis that in long-lived species, pro-longevity genes should be expressed at a higher level than anti-longevity genes, has not been fully examined. Here, we tested this assumption by analyzing the level of LAG expression in the gray whale transcriptome. Nucleotide sequences for all LAGs in the GenAge database (Tacutu et al., 2018) were extracted from RefSeq database (Haft et al., 2018) and BLASTed against the gray whale transcriptome. In total, we used Vsearch (Rognes, Flouri, Nichols, Quince, & Mahe, 2016) to look for 1,422 known LAGs discovered in mouse, fruit fly, and roundworm, of which 601 are pro-longevity and 821 anti-longevity genes. After manual curation and the removal of redundant genes, our analysis shows that 113 (19%) of the transcripts with detectable expression in the gray whale correspond to known pro-longevity genes and only 77 (9%) to anti-longevity genes (Dataset S5).

A noteworthy finding is the lower than expected number of LAG homologues found both for anti- and pro-longevity genes in the gray whale, particularly in view of the assumed high evolutionary conservation of LAGs (Yanai et al., 2017). This, however, has a technical explanation, namely the fact that the gray whale did not have a reference genome until now, and that the transcriptome analysis was performed *de novo*, whereas, for model organisms, there are already numerous well-annotated transcriptomes and genomes. Thus, an incomplete mapping of LAGs is to be expected due to this reason.

In our analysis, in all transcriptomes examined, we found more pro- than anti-longevity genes being expressed. This is despite the fact that, based on the GenAge database (Tacutu et al., 2018) (currently the most comprehensive repository in terms of LAGs) more anti-longevity genes than pro-longevity genes were reported in the literature for each model organism. Several explanations for this finding can be suggested: 1) some anti-longevity genes might not be in the genome entirely because they have been evolutionarily less favored in long-lived species (examples could include: Eef1e1, Trpv1, Pou1f1, all anti-longevity genes that exist in mice, but that do not exist in gray whales); 2) some anti-longevity genes might have a very small expression that is silenced/less detectable in young adults (which is also the case of our gray whale samples) but they might be expressed later in life, a phenomenon is known as antagonistic pleiotropy (an example of a gene with antagonistic pleiotropy characteristics is *prop-1* (Austad & Hoffman, 2018), for more discussions about antagonistic pleiotropy, see (Tacutu et al., 2012; Yanai, Budovsky, Barzilay, Tacutu, & Fraifeld, 2017); 3) some anti-longevity genes might be expressed through the entire lifespan, however at a constantly low level (this case would not appear in our datasets because it is not detectable - so it points to technical difficulty); and finally, 4) some anti-longevity genes might be expressed only on a conditional basis, for example, as co-expression with other genes in response to certain stimuli (again, it is very difficult to detect in sequencing experiments like this one). For the full list of expressed pro- and anti-longevity genes please see Dataset S5.

Two additional points should be stressed in this context: 1) first, as we previously showed, despite high evolutionary distances, the lifespan effects obtained when manipulating orthologous LAGs in different model organisms leads mostly to concordant results (Yanai et al., 2017), and 2) second, the experiments resulting in lifespan extension, and even more so, those where overexpressing LAG results in lifespan extension, are more definite in terms of evaluating the impact of a given gene on longevity than those resulting in lifespan reduction (Yanai et al., 2017). This means that focusing on LAGs from overexpression studies will be more definite and unambiguous. Then, if a pro-longevity LAG was found by overexpression, it is intuitive to expect that this LAG is also highly expressed in longed-lived mammals such as gray whales. Complementarily, when the lifespan-extending effect for a given anti-longevity LAG was found by knock-out or knock-down experiments, its expression in adult gray whales should most likely be at a relatively low level or undetectable. The same could be expected for the overexpression experiments of anti-longevity genes which result in lifespan reduction. The following results of our analysis support these suggestions. Indeed, the vast majority of LAGs found through overexpression experiments, whose orthologs were also identified in the gray whale transcriptome, were pro-longevity (n = 30) and only three were anti-longevity (Table 1). Remarkably, the normalized expression level (TPM) of these 30 pro-longevity genes in the gray whale transcriptome was several-fold higher than the average expression of anti-longevity genes (Fig. 3). Even compared to the average gene expression of the whole gray whale transcriptome, pro-longevity genes displayed 2.6-fold and 4.9-fold higher expression in liver and kidney tissues, respectively (p < 1.37E-6; p < 8.42E-6). Similar results were obtained when comparing the median values of expression (5.8-fold and 6.5-fold increase in liver and kidney, respectively; p < 4.2E-4; 3.77E-5) (Figure 3). In contrast to pro-longevity genes, the three anti-longevity genes showed lower than average expression (68% in liver and 34% in kidney; p < 5E-4) compared to the whole transcriptome. These expression trends were not found in the two mouse transcriptomes that were used in our analysis, even though overall a similar number of LAGs was found (Data not shown).

**Table 1.** Longevity-associated genes (LAGs) from overexpression experiments found in the *de novo* transcriptome of the gray whale. All transcripts from the gray whale transcriptome were aligned to known nematode, fly and mouse LAGs. For comparison, the average TPM for the whole transcriptome is 8.8 for both liver and kidney. Median TPM for the whole transcriptome is 1.2 and 1.7 for liver and kidney, respectively.

| Orthologs from | LAG type | Number of genes | Liver Average (TPM) | Liver Median (TPM) | Kidney Average (TPM) | Kidney Median (TPM) |
| --- | --- | --- | --- | --- | --- | --- |
| Mice | Pro | 13 | 30 | 9 | 67 | 11 |
|  | Anti | 1 | 13.6 | 13.6 | 0.4 | 0.4 |
| Flies | Pro | 12 | 24 | 6 | 33 | 12 |
|  | Anti | 1 | 2.8 | 2.8 | 2.7 | 2.7 |
| Worms | Pro | 5 | 4 | 2 | 5 | 6 |
|  | Anti | 1 | 0.7 | 0.7 | 5.7 | 5.7 |
| Total | Pro | 30 | 23 | 7 | 43 | 11 |
|  | Anti | 3 | 6 | 3 | 3 | 3 |

**Fig. 3.**
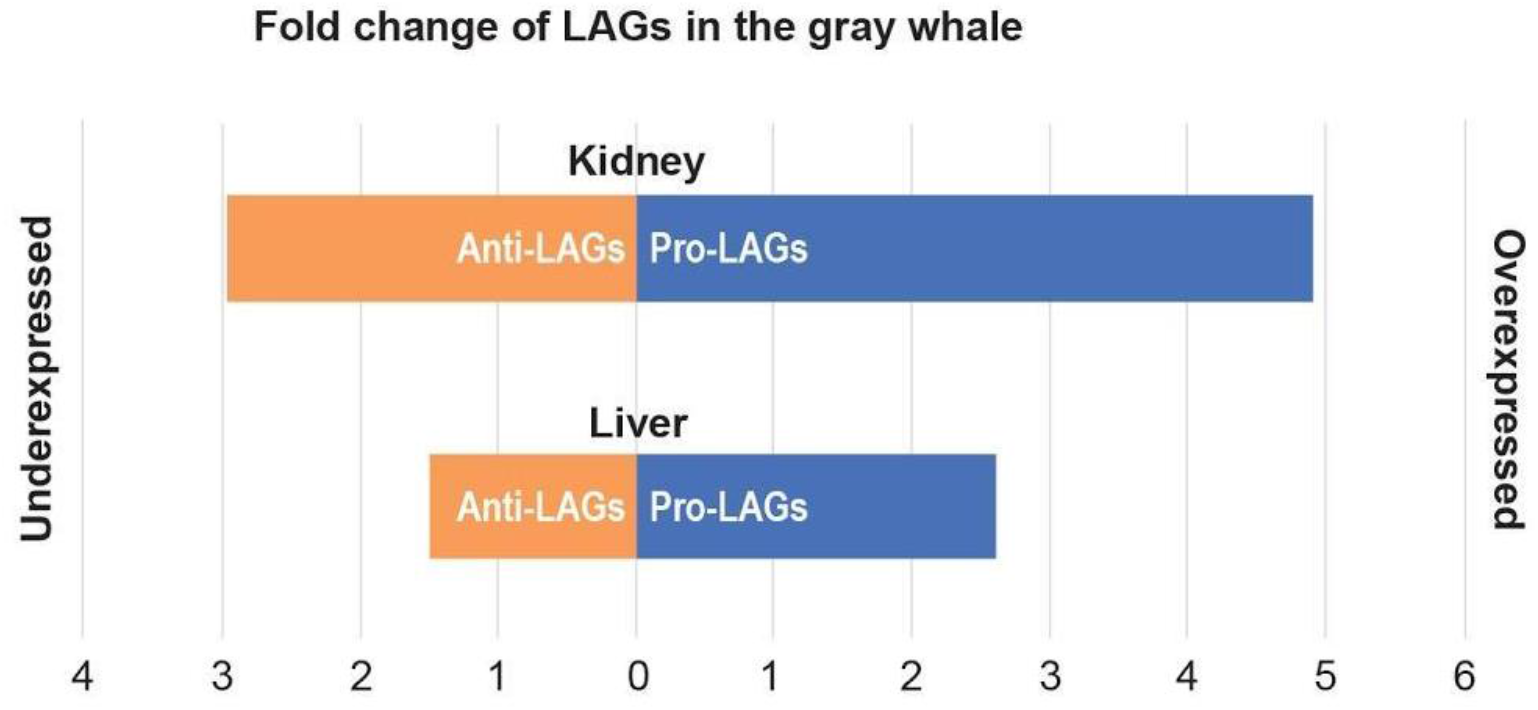
Fold-change in the expression of longevity-associated genes (LAGs) in the *de novo* transcriptome of the gray whale. For this analysis, LAGs were considered only from overexpression experiments in model organisms (*C. elegans*, *D. melanogaster*, *M. musculus*). Displayed are the average fold changes for pro- and anti-longevity genes expressed in the gray whale transcriptome.

As described above, in the gray whale transcriptome, pathways related to DNA repair, autophagy and ubiquitination appear to have increased activity. In regard to this, it was previously suggested that upregulation of stress response genes could result in pro-longevity effects (A. Moskalev et al., 2014). This is further supported by our previous study showing that overexpression of stress-related LAGs results in most cases in lifespan extension (Yanai et al., 2017). Remarkably, in the gray whale transcriptome, a great portion of the pro-longevity LAGs found through overexpression experiments (19 out of 30) are stress-related genes, which display expression values higher than average or median. For example, the heat shock 70kDa protein 1A gene is highly expressed in the gray whale (~25-fold increase, both in liver and kidney), and approximately 118 times higher (p < 5E-4) than in a short-lived species (mouse). Interestingly, the expression of this gene is also high in the naked mole rat (8 fold-increase vs average expression of the whole naked mole rat transcriptome). In contrast, the expression of insulin-like growth factor 1 (IGF-1), an anti-longevity gene, is significantly lower in the gray whale (as well as in other whale species) than in the mouse (18.7 and 10.6-fold decrease in liver and kidney, respectively) (Dataset S5).

If knock-out or knock-down of given LAG results in lifespan extension, it could be expected that its expression in an adult gray whale would be at a relatively low level or undetectable. In line with this expectation, 57 out of the 72 genes (79%) from this group had lower than average expression in liver and 39 out of the 72 genes (54%) had lower than average expression in kidney (Dataset S5). This trend was more obvious for the liver (3.8-fold decrease in gene expression) than for the kidney (1.2-fold decrease). Complementarily, as shown in Table 1, the gray whale orthologous genes of pro-longevity LAGs, whose overexpression led to an increase in the lifespan of model organisms, displayed a markedly higher expression than average in the gray whale transcriptome as well. At the same time, gray whale orthologous genes of pro-longevity LAGs, whose downregulation results in reduced lifespan in model organisms, were found to be expressed at a low level or to be unexpressed at all (Dataset S5). Our approach opens a new avenue for a wide comparative analysis of LAG expression in long- and short-lived species, which could be an important topic for future investigation. The analysis of the gray whale transcriptome and comparison with other mammalian species suggest that the gray whale potentially possesses high resistance to cancer and stress, in part ensuring its longevity.

## Experimental Procedures

### DNA and RNA sampling in tissues

Tissue samples from the kidney (N = 2) and liver (N = 2) were acquired from one adult Gray whale (*Eschrichtius robustus*) female on 31 May 2013, at the seashore Lorino, Chukotka Autonomous Okrug, during the 2013 native Eskimo subsistence harvests, by the indigenous population of Chukotka Autonomous Okrug (at the Mechigmen bay of the Bering Sea, Lorino). The Eskimo have permission to hunt gray whales for food and during one of the hunts, tissue biopsies were taken. No animals were killed specifically for the current study. Sample collection and preparation were previously described in Moskalev et al., 2017(A. A. Moskalev et al., 2017).

### Transcriptome sequencing and Assembly

Total RNA was isolated from frozen tissues using the RNeasy Mini Kit (QIAGEN, Germany) according to the manufacturer’s protocol. RNA quantification was performed on the NanoDrop 1000 (NanoDrop Technologies, USA), and the RNA integrity was assessed using the Agilent 2100 Bioanalyzer (Agilent Technologies, USA). RNA was further treated with DNase I (Thermo Fisher Scientific, USA) and purified using the RNA Clean & Concentrator-5 kit (Zymo Research, USA). The cDNA libraries were prepared using the Illumina TruSeq RNA Sample Preparation Kit v2 (LT protocol) as described in Moskalev et al., 2014 (A. Moskalev et al., 2014). The libraries were sequenced on the Illumina MiSeq System (USA) using the MiSeq Reagent Kit v2 for 500 (2 × 250) cycles. On average, fragment size was 300bp (insertion + adapters), with the insertion size about 180bp. Trimming the samples was performed using Trimmomatic(Bolger, Lohse, & Usadel, 2014). To identify orthologous protein-coding transcripts, we employed a BLAST algorithm against the SwissProt database (Bateman et al., 2017). *De novo* assembly for the four samples was done with the Trinity software stack (Grabherr et al., 2011) and yielded 114,233 contigs. The assembly quality was checked with BUSCO (Simão, Waterhouse, Ioannidis, Kriventseva, & Zdobnov, 2015). Raw data for Brandt’s bat (*Myotis brandtii*) was taken from Series GSE42297 (Seim et al., 2013). Brandt’s bat transcriptome was *de novo* assembled using the RNA Spades assembler, with the default parameters - provided in the GitHub repository (Bushmanova, Antipov, Lapidus, & Prjibelski, 2019).

### Gene Expression Analysis, Filtering, and Expression

To identify orthologous protein-coding transcripts a BLAST algorithm was run against SwissProt (Bateman et al., 2017). Summarization of the results (*de novo* + annotation) was then generated using an in-house built Perl script. The normalization of contig counts was done by computing transcripts per million (TPMs). Genes that were identified and included in subsequent analyses had TPM values of at least 1. To define the high expression threshold for unannotated contigs in the gray whale *de novo* transcriptome, the expression distribution was considered, and an expression level 10x higher than the average of all genes was taken. For a minimal cutoff length for unannotated contigs predicted to be genes, a cutoff of 200bp was used (Bushmanova et al., 2019).

#### Analysis of differential gene expression

Gene ontology (GO) enrichment analysis was carried out in R, using the TopGO package v2.26.0 (Alexa & Rahnenfuhrer, 2016) (http://bioconductor.org/packages/release/bioc/html/topGO.html). Adjusted p-values lower than 0.05 were considered significant. Only reads with one count per million in at least three samples were included in this analysis. For the enrichment analysis of the gray whale transcriptome, the top 100 expressed genes were used. The cutoff was set based on the histogram showing the transcript distribution sorted by their TPM values (Robinson & Oshlack, 2010), in both liver and kidney. The selected 100 top expressed genes account for approximately 50% of the total number of TPMs.

#### Comparisons of transcriptomes

To compare the gene expressions in the gray whale (*Eschrichtius robustus*) across multiple species, we retrieved liver and kidney RNA-Seq data for the bowhead whale (*Balaena mysticetus*), minke whale (*Balaenoptera acutorostrata*), naked mole-rat (*Heterocephalus glaber*), Brandt’s bat (*Myotis brandtii*), human (*Homo sapiens*), cow (*Bos taurus*) and mouse (*Mus musculus*). In this work, only the transcriptome of the gray whale was assembled *de novo*; for all other species previously published transcriptomes were used.

For each species, kidney and liver RNA-Seq assemblies were chosen and quantified with Salmon (Patro, Duggal, Love, Irizarry, & Kingsford, 2017). For mouse and cow, two samples (mouse 1 and 2, cow 1 and 2) were taken in order to investigate variation between samples of the same species. To avoid technical errors - only coding transcripts were taken, and the protein sequences translated from them were aligned. To establish a single point of reference - transcripts were mapped to Uniref90 protein clusters (Suzek, Wang, Huang, McGarvey, & Wu, 2015). For the protein homology search, we used DIAMOND (Buchfink, Xie, & Huson, 2015). The used e-value cutoff was 1E-03. To convert results into ranks - the GO processes from Uniref90 clusters annotations were aggregated by GO biological processes. The protein-coding transcripts, from all transcriptomes, are grouped by GOs and their composite TPMs are computed within each GO term. Terms are ranked based on the composite expression from 1 to 9,779 (number of GO terms in the gray whale’s transcriptome). A phylogenetic tree was built in the TimeTree software (Kumar, Stecher, Suleski, & Hedges, 2017), with default parameters. Solid circles mark nodes that map directly to the NCBI Taxonomy and empty circles indicate nodes that were created during the polytomy resolution process which is described in (Hedges, Marin, Suleski, Paymer, & Kumar, 2015). For each GO category of interest, we randomly generated 10000 pseudo-GO categories with the same number of random Uniref90 clusters.

### LAGs analysis

The comparison with known mouse, fly and worm LAGs was done using data from GenAge database (build 19, 24/06/2017)(Tacutu et al., 2018). Yeast, bacteria and fungus LAGs were excluded from the analysis, resulting in a total of 1168 genes being included in the analysis. Of these, 486 had pro-longevity annotations and 682 had anti-longevity annotations (note: several LAGs are annotated as both pro- and anti-longevity depending on the performed genetic interventions). Nucleotide sequences for these LAGs were exported and aligned using the Vsearch (Rognes et al., 2016) (github.com/torognes/vsearch) sequence alignment tools (parameters are provided inside wdl workflows in the GitHub repository: github.com/antonkulaga/gray-whale-expressions). The results were then manually cleaned to remove duplicates.

### Statistics

For a comparison between different transcriptomes, the Spearman rank correlation was used. The Spearman correlation of two samples is defined as a regular (Pearson’s) correlation between the ranks of values in each sample. To compute the test for significance (*p*-value) using *t* is distributed approximately as Student’s t-distribution with n − 2 degrees of freedom under the null hypothesis (Press, Flannery, Teukolsky, & Vetterling, 1992). A justification for this result relies on a permutation argument(Kendall & Stuart, 1973). For LAGs analysis, we used a hypothesis that the expression of each LAG is described by the Poisson stream of reads. It obeys Poisson distribution with intensity parameter l computed using known TPM values as well as the total number of reads. A cumulative expression is a sum of a large number of Poisson random variables, so it can be approximated using normal distribution according to the Central limit theorem. LAGs expression values have a distribution like a ratio of two Gaussian (normal) variables. We used (numerically computed) CDF of this distribution to get the *p*-values (Hinkley, 1969).

## URLs

This Whole Genome Shotgun project has been deposited at DDBJ/ENA/GenBank under the accession NTJE000000000. Data available at www.ncbi.nlm.nih.gov/nuccore/NTJE000000000 and www.ncbi.nlm.nih.gov/Traces/wgs/?val=NTJE01.

Code availability: github.com/antonkulaga/gray-whale-expressions

## Accession numbers

Bowhead whale: liver tissue SRR1685415, kidney tissue SRR1685390.

Minke whale: liver tissue SRR919296, kidney tissue SRR919295.

Naked mole-rat: liver tissue SRR2123747, kidney tissue SRR2124226.

Human: liver tissue GSM1698568, kidney tissue GSM1698570.

Mouse: liver tissue SM1400574, kidney tissue GSM219518.

Cow: liver tissue GSM1020724, kidney tissue GSM1020723.

Brandt’s bat: liver tissue GSM1037380, GSM1037382, kidney tissue GSM1037381, GSM1037383.

## Acknowledgments

We are very grateful for the help received from Dr. Vered Chalifa-Caspi and Dr. Michal Gordon, from the Bioinformatics Core Facility in the Ben-Gurion University of the Negev. We would also like to thank Dr. Sorel Cahan for his useful comments on the manuscript.

## Contributions

This collaborative study was carried out by the research groups of VEF, AAM and RT. All the samples used in the project have been provided by AAM’s group.

Bioinformatics data processing and analyses of genetic variation data were carried out by DT, AK, MJ. DN performed the statistical analysis. Library construction, sequencing and genome assembly for the draft reference genome were carried out by AVS, AVK and DT. VEF coordinated the research and together with AAM and RT provided supervision for the project. ER provided additional insights into the bioinformatics methodology. VEF, AAM, DT, and RT conceptually designed the experiments and the analyses. DT, RT, and VEF wrote and edited the manuscript. All authors have read and approved the final manuscript.

